# Multi valent DNA vaccine against group A human rotavirus: an *in-silico* investigation

**DOI:** 10.1101/2020.01.13.903781

**Authors:** Kunal Dutta

**Affiliations:** Microbiology and Immunology Laboratory, Department of Human Physiology, Vidyasagar University, West Bengal, Medinipur-721102, India

**Keywords:** group A rotavirus (GARV), VP7, VP8, epitopes, *in silico* cloning, DNA-Vaccine

## Abstract

Gastroenteritis due to single rotavirus causes huge economic loss annually. Severity of rotaviral diarrhoea among children is primarily manifested by different combinations of G and P types. Rotavirus surveillance studies resulted in two ambitious globally licensed vaccine namely, Rotarix and RotaTaq and a few other. However, post-vaccination surveillance studies indicate, vaccine failure and other complications such as intussusception, environmental enteric dysfunction, *etc*. Herein, we design a multivalent DNA vaccine against rotavirus and tested its efficiency by using *in silico* tools. Two main neutralizing rotaviral antigens *i*.*e*, VP7 and VP8 were taken into account and respectively 390, 450 known sequences of different serogroup have been analyzed to obtain a consensus sequence for epitope prediction. Epitopes specific for MHC-I and -II were predicted using IEDB and chosen based on their best IC_50_ value and CPR. A good binding profile with a monoclonal antibody specific for B-cell antigens is displayed by all epitopes they were found to be non-allergenic in the human host. Ethnic specificity of the epitopes is also within acceptable range except for South African and Central American populations. We use pBI-CMV1 bidirectional mammalian expression vector to design the DNA vaccine, where we stapled manually integrated epitopes for VP7 and VP8 at MCS1 and 2 respectively. In conclusion, this study provides a new set of data for a new DNA vaccine against rotavirus.

## Introduction

Rotavirus is a double-stranded RNA (ds-RNA) virus, and the ds-RNA is approximately 670 to 3300 base pairs long which is protected by an inner and outer membrane viral proteins (VPs) ^1^. Inner layer composed of VP1, VP2, VP3, and VP6 ^2^ and based on VP6 antigen, rotavirus is sub-grouped from A to G. Whereas, outer envelope consist of VP7 and a unique spike-like protrusion called, VP4 which is subdivided into four major domains namely, head (VP8), body, stalk, and foot (VP5)^3^. So far, most of the identified neutralizing epitopes are located within VP7, VP8 and VP5 region ^4-6^. Molecular mechanism underlying rotavirus neutralization involve specific IgA, that directly targets VP4 (P-type) and VP7 (G-type) ^7^. In humans, there are 10 G-serotypes and 11 P-genotypes ^8^. Numerous studies have suggested that the severity of rotaviral diarrhea is due to different combinations of G and P types ^9^. As a consequence, rotavirus surveillance studies became meaningful to build vaccine development strategies ^10^. Commonly used two licensed vaccines *i.e*., Rotataq (monovalent), Rotarix (pentavalent) and other vaccines in China, India, and Vietnam have been implemented ^11-14^. However, even after implementation of Rotarix in the national immunization schedule, G1P [8] became dominant genotype (52%) followed by G2P [4]. Conversely, G3P [8] becomes dominant wherein RotaTeq has been used ^15^. However, G2P [4] and G9P [8] were dominated in some states of Brazil ^16^. In Finland, a major reduction of rotavirus gastroenteritis among hospitalized children have been noticed after the advent of Rotataq into the National Immunization Schedule ^17^. Furthermore, in low-middle income counties, annual death due to rotavirus gastroenteritis has decreased dramatically after public availability of Rotarix and RotaTeq ^18, 19^. However, some research cautioned that the vaccine’s effectiveness declines with age ^20^ and post-vaccination surveillance studies suggested rotavirus vaccination associated with a large risk of intussusception ^21, 22^. Moreover, a reduced immunogenic property of the vaccine strain arises a question about effectiveness ^23^. For example, in a recent study rotavirus vaccine failure was associated with a chronic inflammatory intestinal disease called, environmental enteric dysfunction (EED), and EED was present in 80% of infants aged 12 weeks ^24, 25^. Another vaccine surveillance study by a research group, reports rotavirus symptomatic infection among unvaccinated and vaccinated children in Valencia, Spain ^26^.

Conversely, DNA vaccine is a boon of reverse vaccinology that offers great immunogenic protection to an individual. DNA vaccine comprises DNA sequence encoding antigen (s) of interest which can stimulate the host’s immune system ^27^. It has a very good safety profile and they are also well-tolerated in humans ^28^. Further, DNA vaccines can initiate cell-mediated as well as humoral immune reactions ^28^. Besides, large scale manufacturing of DNA vaccines can be cost-effective ^29^. A small peptide molecule that can able to elicit and triggers immune reactions inside host cells is called epitope ^29^. Epitopes from a known protein sequence can be designed by using bioinformatic algorithms. Herein, we use such algorithms to predict epitopes for MHC-I and -II by utilizing a consensus sequence of two main neutralizing viral proteins *i.e*., VP7 and VP8. Immunogenic peptides were inserted into a bidirectional mammalian expression vector; pBI-CMV1 which can simultaneously express two DNA inserts from multiple cloning site (MCS) 1 and 2. CpG motif optimization was performed prior to insertion of the immunogenic peptides into the pBI-CMV1.

## Results and Discussion

Public health is a fundamental criterion to build a disease-free healthy nation ^30^. A vaccine of a particular disease provide great immunogenic protection to an individual in his / her complete life and it also consecutively reduces personal healthcare cost^31^. Moreover, the healthcare economy is indirectly reflected in the annual healthcare cost burden of a nation, which is also immensely reduced by implementation of vaccines in the national immunization schedule ^32^. In developed countries and also countries from the global south, childhood diarrhoea due to single rotavirus causes huge economic loss ^33^. Epidemiological pre-vaccination surveillance studies on rotavirus help to build vaccine development strategies that resulted in two ambitious globally licensed vaccines. However, rotavirus vaccines made from conventional vaccinology (*e.g*. attenuated or live-attenuated) arises questions about their performance ^20-22, 24^. Conversely, reverse vaccinology becomes boon to experimental scientists before the formulation of a new vaccine candidate ^34^. DNA vaccines are also safe and they present intrinsic adjuvant properties, including cell-mediated immune response, and can result in long-lasting immunity ^35^. Phase-I trials of DNA vaccine against the Ebola virus and chimeric HIV vaccine have already been initiated by the NIH research program ^36^. Herein, we computationally designed a multivalent DNA vaccine against group A human rotavirus. The inherent DNA stability and ability to elicit cell-mediated immune responses (CD4+ and CD8+ T cells), armament DNA vaccines for rapidly responding infectious diseases ^37^. Further, needle-free applications and low maintenance offer easy transport and availability ^38^. Several reports from pre-vaccination to post-vaccination period suggested that, the main rotavirus neutralizing antigens were from VP7 and VP8 (Figure 1). Therefore, we have analyzed 398 and 358 known sequences of different serogroup respectively for VP7 and VP8. On average Shanon variability index value was found to be 1, Shimshon variability index value was found to be > 0.6 and the Wu-Kabat variability was about 10, which collectively indicates variations in the amino acid sequence of the VP7 among different genogroup. Conversely, the average Shanon, Simpson, and Wu-Kabat variability index value for VP8 was comparatively lower than VP7. Phylogenetic tree of VP7 suggests rotavirus strains of the same genotypes fall within a common leaf with only a few exceptions (Figure 1). Conversely, phylogenetic tree of VP8 displayed multidirectional evolution, where rotavirus strains of the same genotypes appear on different branches of the same tree (Figure 2). Moreover, the pair-wise distance matrix for VP7 and VP8 are also in agreement with the data obtained from the phylogenetic trees (Figure 4). As mentioned early, epitopes were selected based on their best IC_50_ value and CPR and the rest are summarised in Table S1. Epitopes for both MHC classes for VP7 and VP8 with their best binding free energies were presented in Figure 6 along with their interacting amino acid residues. Majority of epitopes have an immunogenic propensity of > 1, indicating predicted epitopes can elicit an adequate immune response. Besides, all epitopes were found to be non-allergenic to human host meaning safe for human use (Table S2). The high polymorphic nature of the Human Leucocyte Antigen (HLA) system, confer their ethnic specificity, which leads to poor immunogenicity of a particular epitope to a particular population ^39^ and may cause an optimal immune response to other population. Bi-directional mammalian vector pBI-CMV1 simultaneously express VP7 and VP8 distinctive immunogenic peptides for both MHC classes (Figure 5).

**Figure 1.**
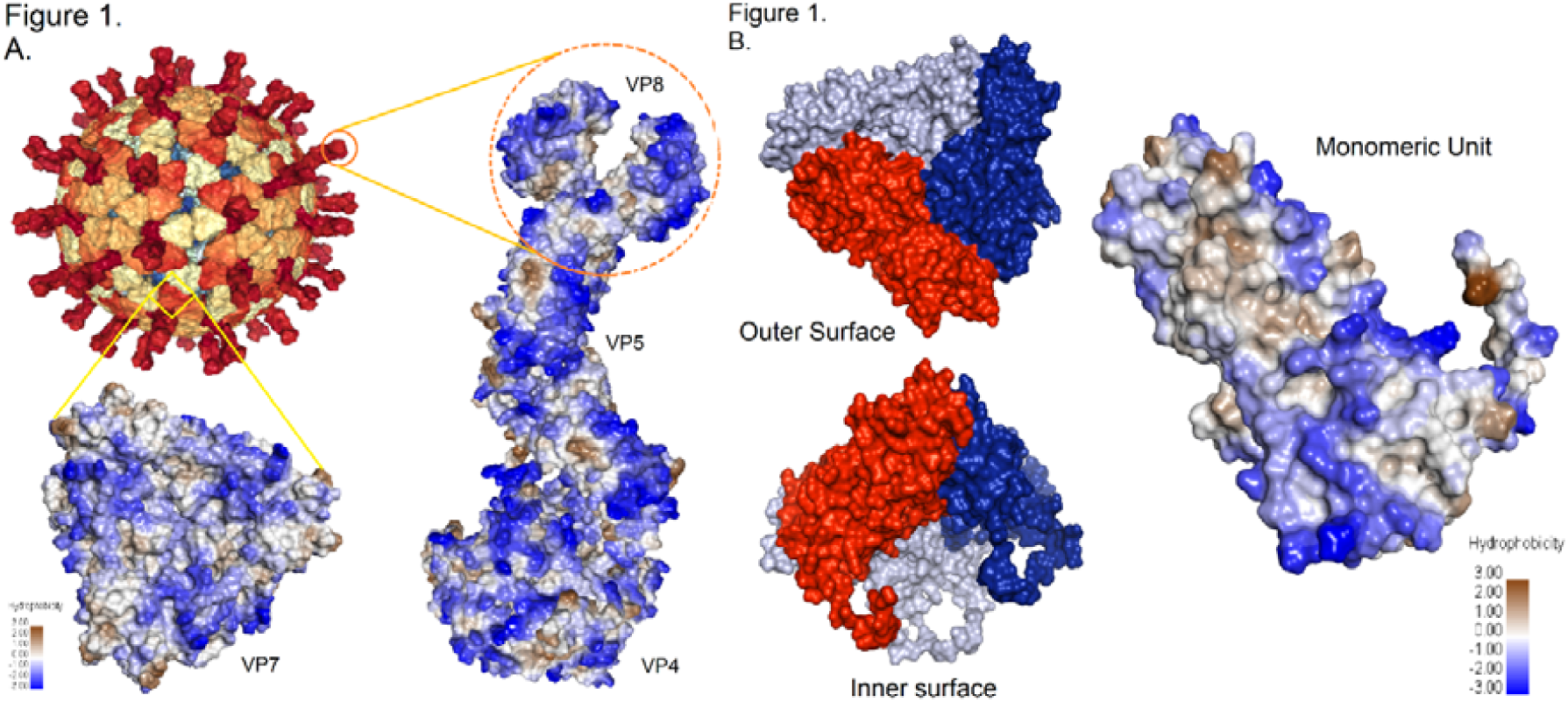
Anatomy of two main neutralizing rotavirus outer membrane protein structure. (A.) Viral protein 7 and anatomy of viral protein 8. (B.) Anatomy of viral protein 7. Color scaled is based on the hydrophobicity index of the monomeric of VP7.

**Figure 2.**
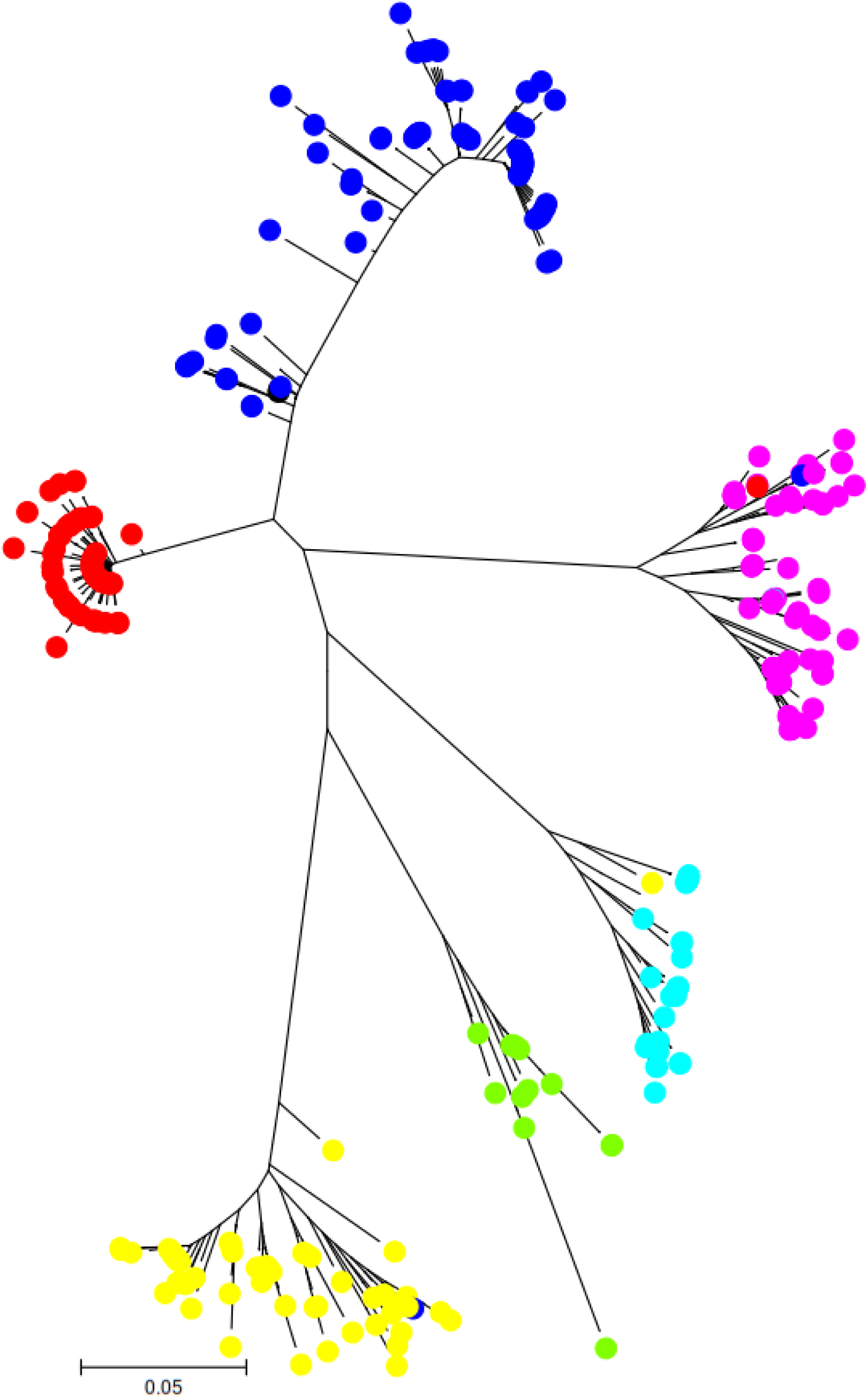
Phylogenetic tree of viral protein 7. The evolutionary history was inferred using the Minimum Evolution method. The optimal tree with the sum of branch length = 3.53915720 is shown. The tree is drawn to scale, with branch lengths in the same units as those of the evolutionary distances used to infer the phylogenetic tree. The analysis involved 398 amino acid sequences. All positions containing gaps and missing data were eliminated. There was a total of 84 positions in the final dataset. The tree is drawn using 1000 bootstrap replication. Red (G9), blue (G3), purple (G1 [P8]), cyan (G8 [P6], G1), light green (G10 [P8]) and yellow (G2 [P4]).

**Figure 3.**
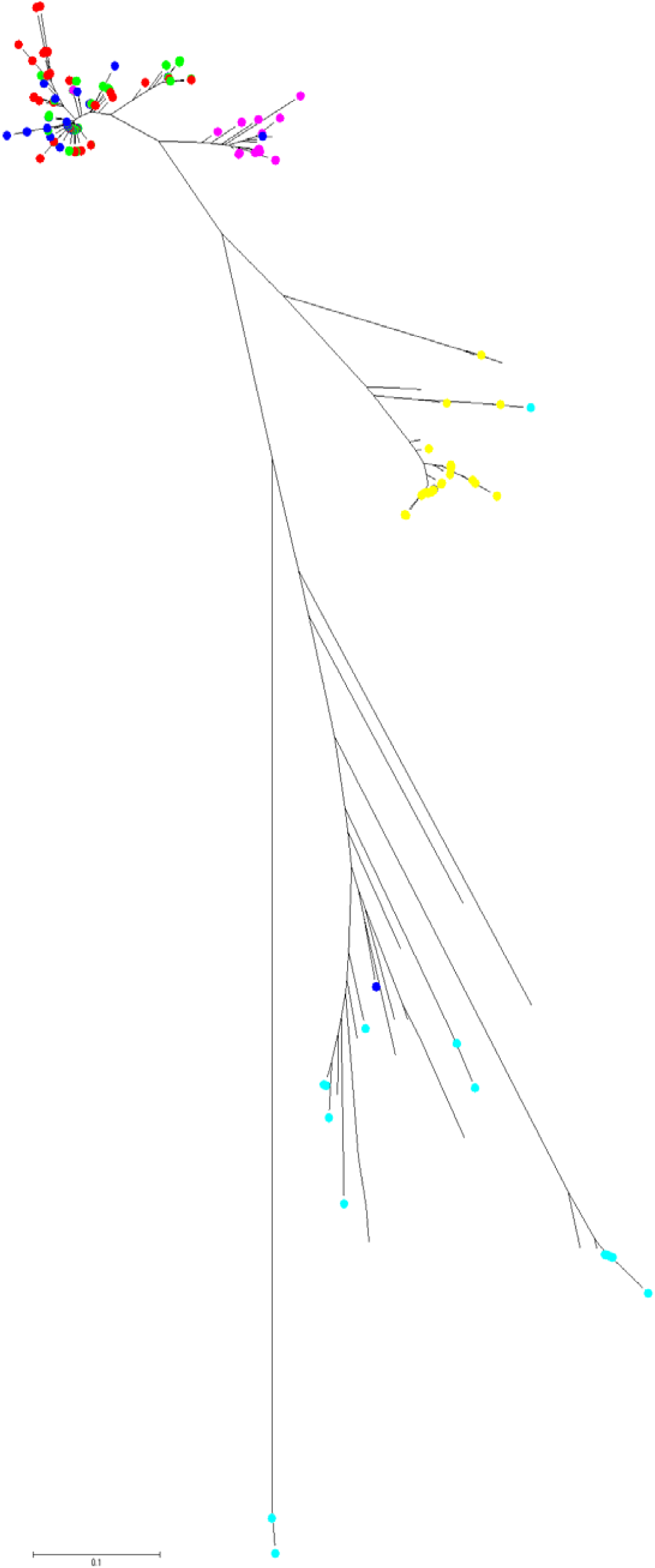
Phylogenetic tree of viral protein 8. The evolutionary history was inferred using the Minimum Evolution method. The optimal tree with the sum of branch length = 5.85019170 is shown. The tree is drawn to scale, with branch lengths in the same units as those of the evolutionary distances used to infer the phylogenetic tree. The analysis involved 358 amino acid sequences. All positions containing gaps and missing data were eliminated. There was a total of 64 positions in the final dataset. The tree is drawn using 1000 bootstrap replication. Yellow (G9 [P6]), purple (G2 [P4]/ [P6]/[P8]), blue (G9[P8]/[P2], cyan (G3 [P9], red (G3 [P8]/ [P6]), light green (G1 [P8]).

**Figure 4.**
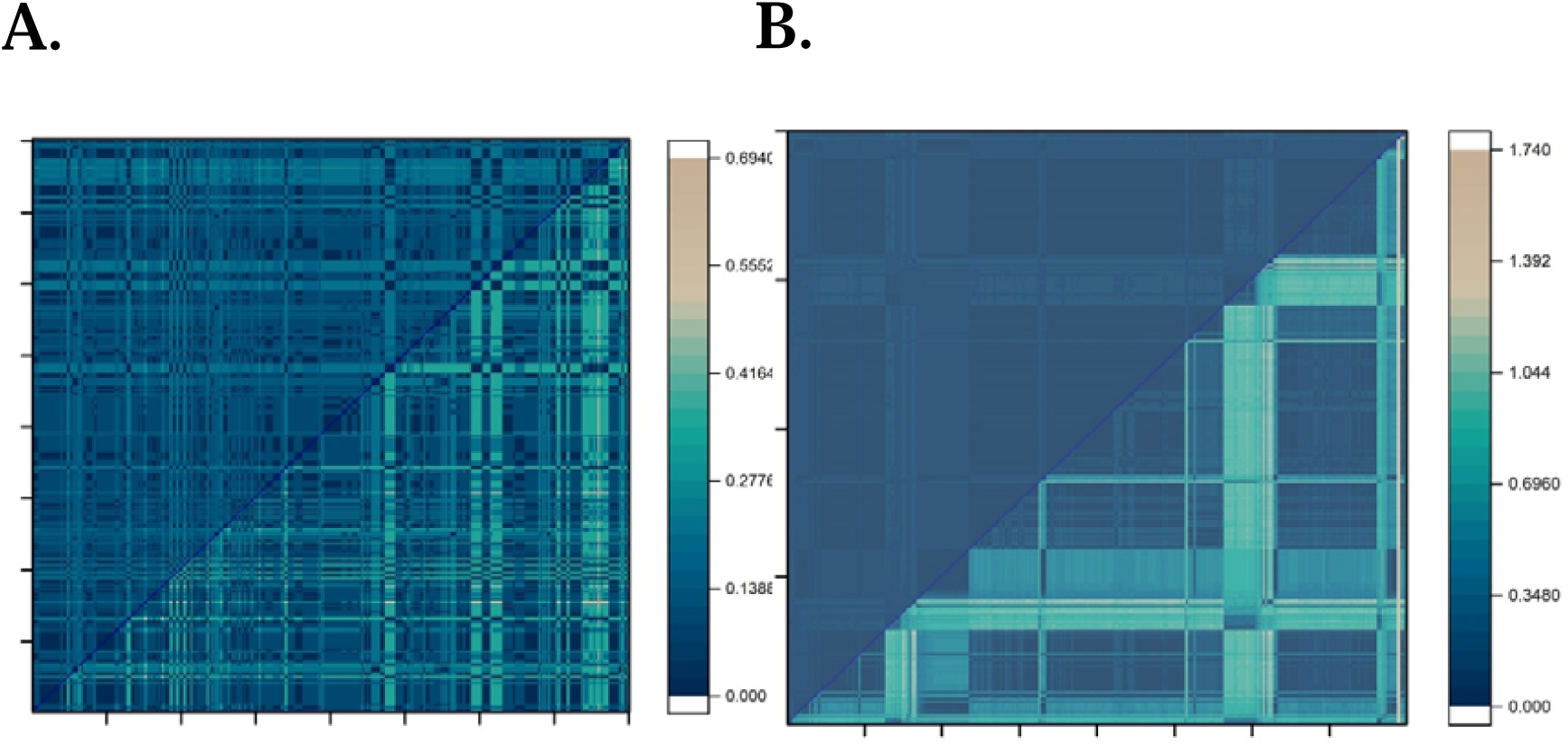
Graphical presentation of the pair-wise distant matrix of viral outer capsid proteins. (A.) Pair-wise distance matrix of viral protein 7 of rotavirus. (B.) viral protein 8 of rotavirus. Color-coded scale at the right side indicates the minimum and maximum distance for the analyzed sequences.

**Figure 5.**
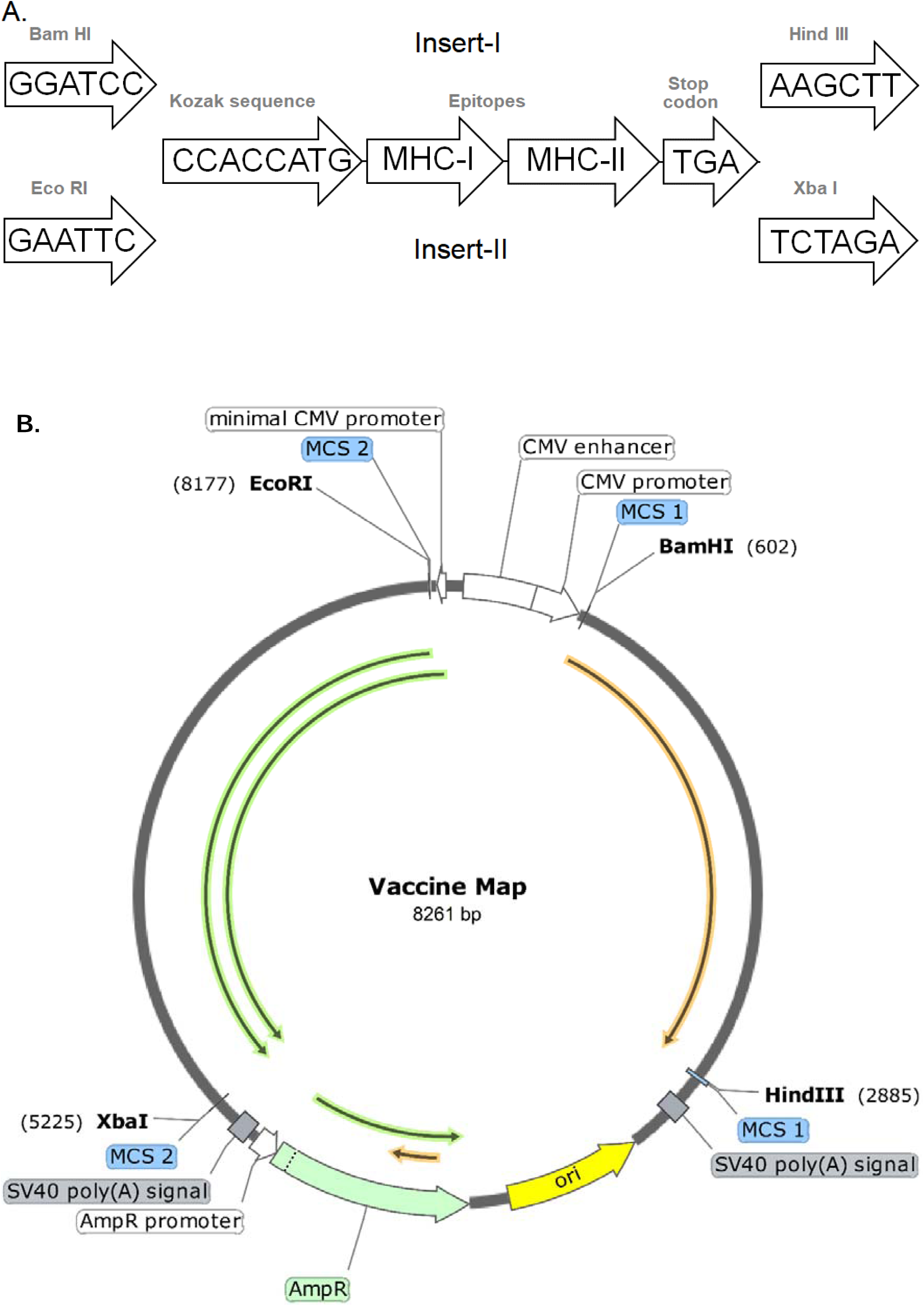
Major steps in the construction of a Rotavirus DNA vaccine. (A.) Functional annotation of the immunogenic insert-I (for VP7) and insert-II (for VP8) with respective type-II restriction endonucleases. (B.) DNA vaccine map against rotavirus with its main features.

**Figure 6.**
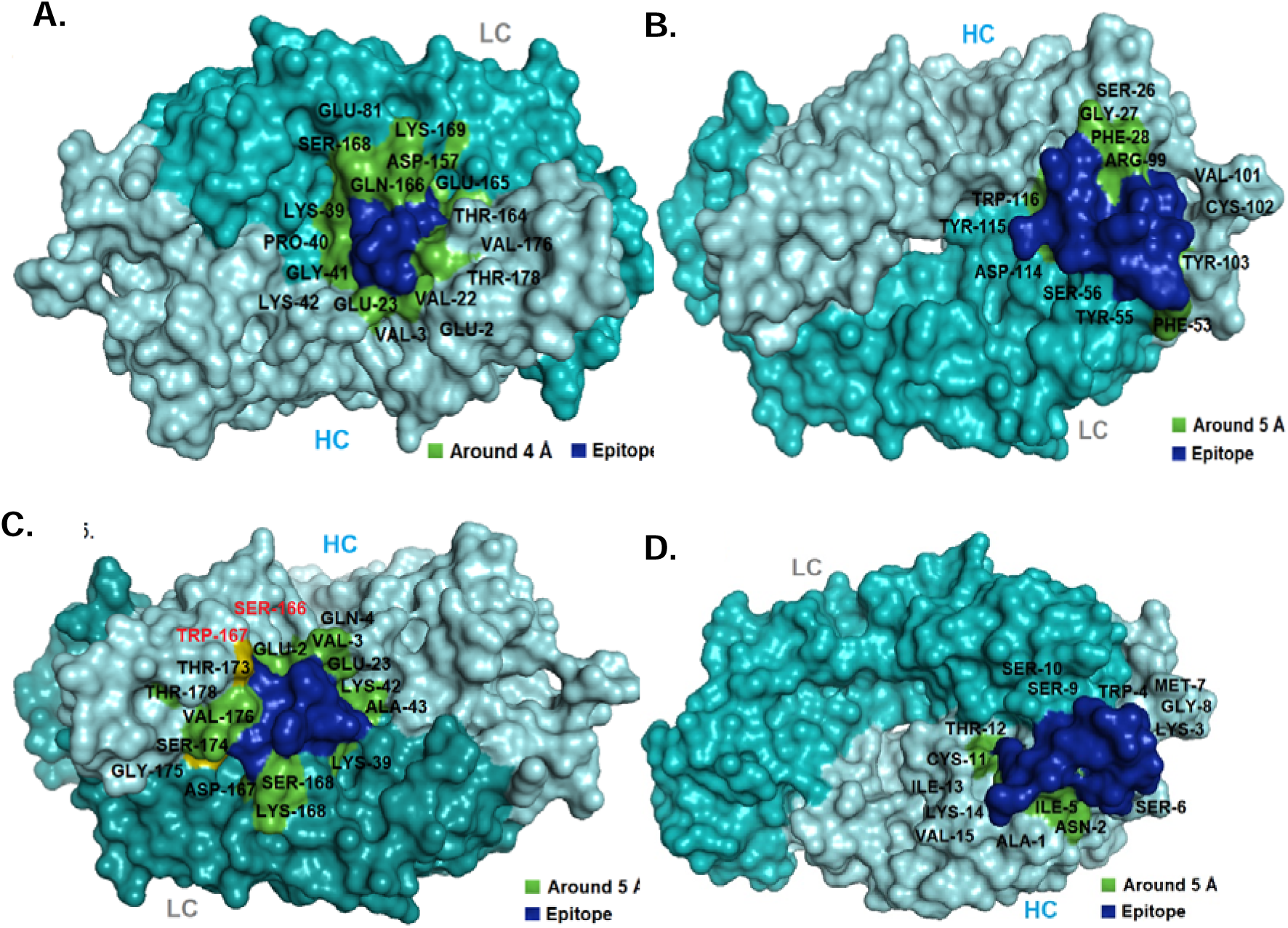
Docking topology of epitopes with Fab of a human monoclonal antibody (mAb) specific for B-cell antigen. (A.) Interaction of GPRENVAVI (VP7-MHC-I). (B.) TSTLCLYYPTEAATE (VP7-MHC-II). (C.) TAFCDFYII (VP8-MHC-I) and (D.) APTAAGVVVEGTNNT (VP8-MHC-II). HC = Heavy chain (cyan), LC = Light chain (gray). Blue color indicates bound epitope and green color indicates interacting amino acid residues in the vicinity.

Therefore, this DNA vaccine will overcome the shortcomings of small population coverage offered by VP7 immunogenic peptide for MHC-II (Figure 7). However, small population coverage for South African and Central American populations (Table S3) can be bypassed by arranging a special type of immune regimen or by designing allele-specific epitope upon needs. Overall, this approach is well suited for constructing a new DNA vaccine for rotavirus.

**Figure 7.**
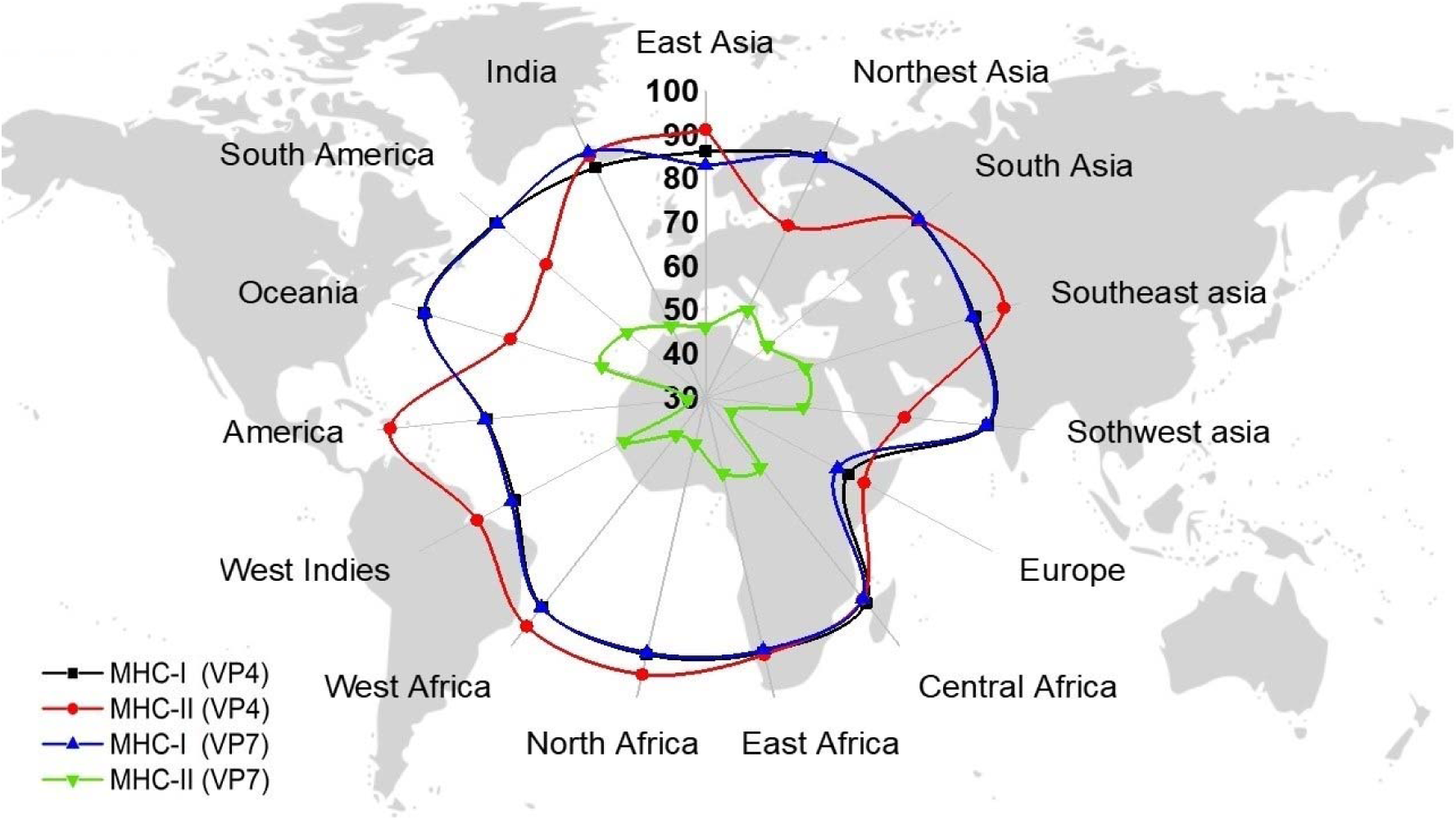
Population coverage offered by predicted epitopes for group A human rotavirus.

## Supplementary files

Supplementary files include Table S1, S2 and S3.

## Acknowledgement

Council of Scientific and Industrial Research (CSIR), Govt. of India, New Delhi, India is also sincerely acknowledged by K.D. for Senior Research Fellowship (SRF), sanction letter no. 09/599 (0082) 2K19 EMR-Z.

## References

1. Estes, M. & Kapikian, A. Fields virology. Philadelphia, PA: Lippencott, Williams and Wilkins (2007).

2. Patton, J., Silvestri, L., Tortorici, M., Vasquez-Del Carpio, R. & Taraporewala, Z. in Reoviruses: Entry, Assembly and Morphogenesis 169–187 (Springer, 2006).

3. Dormitzer, P.R., Nason, E.B., Prasad, B.V. & Harrison, S.C. Structural rearrangements in the membrane penetration protein of a non-enveloped virus. Nature 430, 1053 (2004).

4. Feng, N. et al. Human VP8* mAbs neutralize rotavirus selectively in human intestinal epithelial cells. The Journal of clinical investigation 129 (2019).

5. Nair, N. et al. VP4-and VP7-specific antibodies mediate heterotypic immunity to rotavirus in humans. Science translational medicine 9, eaam5434 (2017).

6. Denisova, E. et al. Rotavirus capsid protein VP5* permeabilizes membranes. Journal of virology 73, 3147–3153 (1999).

7. Liu, K., Yang, X., Wu, Y. & Li, J. Rotavirus strategies to evade host antiviral innate immunity. Immunology letters 127, 13–18 (2009).

8. Gentsch, J.R. et al. Serotype diversity and reassortment between human and animal rotavirus strains: implications for rotavirus vaccine programs. Journal of Infectious Diseases 192, S146–S159 (2005).

9. Sudarmo, S.M. et al. Genotyping and clinical factors in pediatric diarrhea caused by rotaviruses: one-year surveillance in Surabaya, Indonesia. Gut pathogens 7, 3 (2015).

10. Glass, R.I. et al. Rotavirus vaccines: current prospects and future challenges. The Lancet 368, 323–332 (2006).

11. Dennehy, P.H. Rotavirus vaccines: an overview. Clinical microbiology reviews 21, 198–208 (2008).

12. O’Ryan, M. Rotarix™ (RIX4414): an oral human rotavirus vaccine. Expert review of vaccines 6, 11–19 (2007).

13. McCarthy, M. Project seeks to “fast track” rotavirus vaccine. The Lancet 361, 582 (2003).

14. Naik, S.P. et al. Stability of heat stable, live attenuated Rotavirus vaccine (ROTASIIL®). Vaccine 35, 2962–2969 (2017).

15. Kirkwood, C.D., Boniface, K., Barnes, G.L. & Bishop, R.F. Distribution of rotavirus genotypes after introduction of rotavirus vaccines, Rotarix® and RotaTeq®, into the National Immunization Program of Australia. The Pediatric infectious disease journal 30, S48–S53 (2011).

16. Carvalho-Costa, F.A. et al. Rotavirus genotype distribution after vaccine introduction, Rio de Janeiro, Brazil. Emerging infectious diseases 15, 95 (2009).

17. Hemming, M. et al. Major reduction of rotavirus, but not norovirus, gastroenteritis in children seen in hospital after the introduction of RotaTeq vaccine into the National Immunization Programme in Finland. European journal of pediatrics 172, 739–746 (2013).

18. Desselberger, U. Rotaviruses. Virus research 190, 75–96 (2014).

19. Desselberger, U. & Huppertz, H.-I. Immune responses to rotavirus infection and vaccination and associated correlates of protection. Journal of Infectious Diseases 203, 188–195 (2011).

20. Correia, J.B. et al. Effectiveness of monovalent rotavirus vaccine (Rotarix) against severe diarrhea caused by serotypically unrelated G2P [4] strains in Brazil. The Journal of infectious diseases 201, 363–369 (2010).

21. Esposito, S., Pugni, L., Mosca, F. & Principi, N. Rotarix® and RotaTeq® administration to preterm infants in the neonatal intensive care unit: Review of available evidence. Vaccine 36, 5430–5434 (2018).

22. Glass, R.I., Jiang, B. & Parashar, U. The future control of rotavirus disease: Can live oral vaccines alone solve the rotavirus problem? Vaccine 36, 2233–2236 (2018).

23. Zeller, M. et al. Genetic analyses reveal differences in the VP7 and VP4 antigenic epitopes between human rotaviruses circulating in Belgium and rotaviruses in Rotarix and RotaTeq. Journal of clinical microbiology 50, 966–976 (2012).

24. Ramakrishnan, G., Ma, J.Z., Haque, R. & Petri, W.A. Rotavirus vaccine protection in low-income and middle-income countries. The Lancet Infectious Diseases (2019).

25. Naylor, C. et al. Environmental enteropathy, oral vaccine failure and growth faltering in infants in Bangladesh. EBioMedicine 2, 1759–1766 (2015).

26. Pérez-Ortín, R. et al. Rotavirus symptomatic infection among unvaccinated and vaccinated children in Valencia, Spain. BMC infectious diseases 19, 998 (2019).

27. Donnelly, J.J., Wahren, B. & Liu, M.A. DNA vaccines: progress and challenges. The Journal of Immunology 175, 633–639 (2005).

28. Cann, A.J. in Principles of Molecular Virology (Sixth Edition). (ed. A.J. Cann) 173–220 (Academic Press, Boston; 2016).

29. Zhang, N.-Z. et al. Protective efficacy of two novel DNA vaccines expressing Toxoplasma gondii rhomboid 4 and rhomboid 5 proteins against acute and chronic toxoplasmosis in mice. Expert review of vaccines 14, 1289–1297 (2015).

30. Modi, N., Clark, H., Wolfe, I., Costello, A. & Budge, H. A healthy nation: strengthening child health research in the UK. The Lancet 381, 73–87 (2013).

31. Schneider, J. et al. Enhanced immunogenicity for CD8+ T cell induction and complete protective efficacy of malaria DNA vaccination by boosting with modified vaccinia virus Ankara. Nature medicine 4, 397 (1998).

32. John, J. et al. Rotavirus gastroenteritis in India, 2011–2013: revised estimates of disease burden and potential impact of vaccines. Vaccine 32, A5–A9 (2014).

33. Jin, H. et al. Hospital-based study of the economic burden associated with rotavirus diarrhea in eastern China. Vaccine 29, 7801–7806 (2011).

34. Rappuoli, R. Reverse vaccinology, a genome-based approach to vaccine development. Vaccine 19, 2688–2691 (2001).

35. Nascimento, I. & Leite, L. Recombinant vaccines and the development of new vaccine strategies. Brazilian journal of medical and biological research 45, 1102–1111 (2012).

36. Lambe, T., Bowyer, G. & Ewer, K.J. A review of phase I trials of Ebola virus vaccines: what can we learn from the race to develop novel vaccines? Philosophical Transactions of the Royal Society B: Biological Sciences 372, 20160295 (2017).

37. Panagioti, E., Klenerman, P., Lee, L.N., Van Der Burg, S.H. & Arens, R. Features of effective T cell-inducing vaccines against chronic viral infections. Frontiers in immunology 9, 276 (2018).

38. Ferraro, B. et al. Clinical applications of DNA vaccines: current progress. Clinical infectious diseases 53, 296–302 (2011).

39. Alexander, J. et al. Universal influenza DNA vaccine encoding conserved CD4+ T cell epitopes protects against lethal viral challenge in HLA-DR transgenic mice. Vaccine 28, 664–672 (2010).

