## Supplementary files for "Multi valent DNA vaccine against group A human rotavirus: an *in-silico* investigation"

Table S1. Properties of the best predicted epitopes

| Sl. No. | Epitopes | Molecular weight | Isoelectric point | Solubility | $T_{1/2}$ (h) | Kcal $Mol^{-1}$ | | |
|---|---|---|---|---|---|---|---|---|
| | | | | | | Initial Energy (sOPEP energy) | Minimum conformational energy | Docking energy |
| 1 | LTSTLCLYY | 1076.32 | 5.11 | - | 5.5 | -11.52 | -7.74 | -27.38 |
| 2 | SLNSAAFYY | 1035.11 | 5.24 | - | 1.9 | -11.01 | -7.85 | -39.91 |
| 3 | YTDIASFSV | 1002.09 | 3.80 | - | | -8.10 | -8.81 | -40.50 |
| 4 | NWKKWWQVF | 1321.55 | 10.0 | - | 1.4 | -16.37 | -9.07 | -33.73 |
| 5 | VLMKYDATL | 1053.28 | 5.81 | - | 100 | -10.79 | -8.02 | -30.06 |
| 6 | MMRINWKKW | 1292.63 | 11.17 | + | 30 | -11.25 | -7.62 | -39.85 |
| 7 | TVVDYVNQI | 1050.18 | 3.80 | - | 7.2 | -9.89 | -8.03 | -39.14 |
| 8 | AMSKRSRSL | 1035.23 | 12.01 | + | 4.4 | -8.95 | -7.61 | -39.88 |
| 9 | KEYTDIASF | 1073.15 | 4.37 | + | 1.3 | -9.47 | -7.85 | -42.16 |
| 10 | YVNQIIQAM | 1079.28 | 5.52 | - | 2.8 | -11.68 | -6.96 | -30.25 |
| 11 | WKDTLSQLF | 1137.30 | 5.84 | - | 2.8 | -13.03 | -7.58 | -20.71 |
| 12 | LYCDYNVVL | 1101.28 | 3.80 | - | 5.5 | -5.88 | -6.94 | -44.92 |
| 13 | VATAEKLVI | 943.14 | 5.97 | + | 100 | -11.69 | -7.98 | -36.13 |
| 14 | FLTSTLCLY | 1060.27 | 5.62 | - | 1.1 | -11.34 | -9.07 | -39.33 |
| 15 | RENVAVIQV | 1027.18 | 6.00 | + | 1.0 | -3.75 | -1.07 | -5.18 |
| 16 | DVVDGVNHK | 982.06 | 5.21 | + | 1.1 | -7.20 | -6.41 | -45.14 |
| 17 | GPRENVAVI | 954.09 | 6.00 | + | 30 | -7.77 | -6.82 | -52.89 |
| 18 | RSLNSAAFY | 1028.12 | 8.75 | - | 1.0 | -10.38 | -7.97 | -34.95 |
| 19 | MKYDATLQL | 1082.28 | 5.59 | + | 30 | -10.53 | -7.78 | -43.01 |
| 20 | KWWQVFYTV | 1256.45 | 8.59 | - | 1.3 | -14.91 | -7.62 | -51.85 |
| 21 | RMMRINWKK | 1262.6 | 12.02 | + | 1.0 | -9.10 | -7.72 | -30.76 |
| 22 | RSRSLNSAA | 961.04 | 12.0 | + | 1.0 | -9.28 | -8.61 | -36.41 |
| 23 | NPMDITLYY | 1129.29 | 3.80 | - | 1.4 | -5.16 | -1.22 | -44.80 |

| | | | | | | | | |
|---|---|---|---|---|---|---|---|---|
| 24 | ANKWISMGSSCTIKV | 1624.93 | 9.32 | - | 4.4 | -14.23 | -1.33 | -21.26 |
| 25 | DATLQLDMSELADLI | 1647.86 | 3.32 | + | 1.1 | -18.83 | -1.14 | -38.67 |
| 26 | SVYFKEYTDIASFSV | 1755.94 | 4.37 | - | 1.9 | -16.62 | -1.49 | -72.23 |
| 27 | KKWWQVFYTVVDYVN | 1975.28 | 8.43 | - | 1.3 | -27.57 | -1.45 | 49.87 |
| 28 | SCTIKVCPLNTQTLG | 1577.87 | 7.79 | - | 1.9 | -6.89 | -1.58 | -51.46 |
| 29 | VATAEKLVITDVVDG | 1529.75 | 4.03 | + | 100 | -13.68 | -1.18 | -29.17 |
| 30 | FYTVVDYVNQIIQAM | 1804.07 | 3.80 | - | 1.1 | -23.10 | -1.34 | -45.76 |
| 31 | INWKKWWQVFYTVVD | 2012.31 | 8.50 | - | 20 | -30.29 | -1.31 | -71.58 |
| 32 | NVAVIQVGGSDVLDI | 1498.68 | 3.56 | - | 1.4 | -17.29 | -1.40 | -34.78 |
| 33 | CDYNVVLMKYDATLQ | 1776.05 | 4.21 | - | 1.2 | -10.53 | -2.53 | -38.96 |
| 34 | DYVNQIIQAMSKRSR | 1809.06 | 9.99 | + | 1.1 | -22.04 | -1.38 | -21.67 |
| 35 | WQVFYTVVDYVNQII | 1887.17 | 3.80 | - | 2.8 | -25.02 | -1.02 | -48.05 |
| 36 | YVNQIIQAMSKRSRS | 1781.06 | 11.00 | + | 2.8 | -22.54 | -1.43 | -51.49 |
| 37 | TSTLCLYYPTEAATE | 1662.83 | 3.79 | - | 7.2 | -13.17 | -1.14 | -52.89 |
| 38 | EETFLTSTLCLYYPT | 1781.01 | 3.80 | - | 1.0 | -17.04 | -1.60 | -47.61 |
| 39 | PQTERMMRINWKKWW | 2090.48 | 11 | + | 20 | -29.65 | -1.47 | -21.08 |
| 40 | PMDITLYYYQQTDEA | 1851.01 | 3.49 | - | 20 | -9.16 | -1.85 | -20.29 |
| 41 | TDEANKWISMGSSCT | 1629.77 | 4.37 | + | 7.2 | -18.85 | -1.58 | -25.91 |
| 42 | DYNVVLMKYDATLQL | 1786.06 | 4.21 | - | 1.1 | -13.28 | -2.29 | -37.37 |
| 43 | WWQVFYTVVDYVNQI | 1960.22 | 3.80 | - | 2.8 | -27.26 | -1.09 | -27.40 |
| 44 | TTATCTIRNCKKLGP | 1606.92 | 9.50 | + | 7.2 | -12.04 | -1.20 | -33.02 |
| 45 | YNVVLMKYDATLQLD | 1786.06 | 4.21 | - | 2.8 | -11.52 | -2.74 | -29.65 |
| 46 | VVDYVNQIIQAMSKR | 1764.07 | 8.56 | + | 100 | -22.98 | -1.35 | -34.72 |
| 47 | INDNSWKDTLSQLFL | 1793.97 | 4.21 | - | 20 | -21.14 | -1.35 | -52.01 |
| 48 | MTAFCDFYI | 1110.31 | 3.80 | - | 30 | -12.93 | -7.99 | -44.83 |
| 49 | LLAPTAAGV | 811.97 | 5.52 | - | 5.5 | -6.80 | -7.95 | -35.43 |
| 50 | TTFNPPVDY | 1053.12 | 3.80 | - | 7.2 | -3.85 | -7.70 | -32.47 |
| 51 | SETRSYTLF | 1103.18 | 5.72 | + | 1.9 | -3.49 | -1.24 | -35.93 |
| 52 | TPNVTTKYY | 1086.21 | 8.17 | - | 7.2 | -3.40 | -7.61 | -36.59 |
| 53 | TQNGSYSQY | 1047.03 | 5.18 | - | 7.2 | -3.73 | -1.01 | -37.02 |
| 54 | PPVDYWMLL | 1133.37 | 3.80 | - | 20 | -13.91 | -7.78 | -52.00 |
| 55 | ETRSYTLFG | 1073.16 | 6.10 | + | 1.0 | -4.54 | -8.22 | -36.20 |

| | | | | | | | | |
|---|---|---|---|---|---|---|---|---|
| 56 | YWMLLAPTA | 1065.30 | 5.52 | - | 2.8 | -11.44 | -6.68 | -41.74 |
| 57 | GSYSQYGPL | 971.02 | 5.52 | - | 30 | -5.45 | -6.59 | -41.85 |
| 58 | QSTPKLYAV | 1006.15 | 8.59 | - | 0.8 | -8.00 | -8.02 | -42.81 |
| 59 | NMTAFCDFY | 1111.25 | 3.80 | - | 1.4 | -11.66 | -9.87 | -41.11 |
| 60 | DRWLATILV | 1086.30 | 5.84 | - | 1.1 | -14.24 | -7.87 | -38.55 |
| 61 | TAFCDFYII | 1092.26 | 3.80 | - | 7.2 | -11.06 | -7.94 | -56.34 |
| 62 | TSETRSYTL | 1057.13 | 5.66 | + | 7.2 | -3.85 | -1.12 | -52.58 |
| 63 | TQNGSYSQY | 1047.03 | 5.18 | - | 7.2 | -3.75 | -1.01 | -37.02 |
| 64 | APTAAGVVV | 783.92 | 5.57 | - | 4.4 | -5.84 | -5.74 | -30.18 |
| 65 | TTFNPPVDY | 1053.12 | 3.80 | - | 7.2 | -3.85 | -7.73 | -32.47 |
| 66 | ETPNVTTKY | 1052.14 | 6.10 | + | 1 | -3.19 | -7.71 | -30.49 |
| 67 | PVDYWMLLAPTAAGV | 1603.88 | 3.80 | - | 20 | -22.22 | -1.03 | -30.62 |
| 68 | TSETRSYTLFGTQEQ | 1747.81 | 4.53 | + | 7.2 | -5.12 | -1.41 | -39.22 |
| 79 | STTNYDSVNMTAFCD | 1688.76 | 3.56 | - | 1.9 | -11.35 | -1.47 | -24.83 |
| 70 | QSTPKLYAVMKHNGK | 1702.00 | 10 | + | 0.8 | -18.65 | -1.58 | -38.14 |
| 71 | YSQYGPLQSTPKLYA | 1715.19 | 8.43 | - | 2.8 | -12.47 | -1.15 | -49.93 |
| 72 | NASQTQWKFIDVVKT | 1764.97 | 8.59 | + | 1.4 | -20.84 | -1.15 | -26.87 |
| 73 | NYDSVNMTAFCDFYI | 1802.99 | 3.56 | - | 1.4 | -17.36 | -1.45 | -42.10 |
| 74 | KLYAVMKHNGKIYTY | 1829.19 | 9.70 | - | 1.3 | -20.31 | -1.00 | -36.29 |
| 75 | LATILVEPNVTSETR | 1642.85 | 4.53 | + | 5.5 | -10.63 | -9.88 | -39.76 |
| 76 | PKLYAVMKHNGKIYT | 1763.13 | 9.84 | - | 20 | -20.90 | -1.18 | -33.70 |
| 77 | MTAFCDFYIIPREEE | 1864.11 | 4.00 | + | 30 | -15.46 | -1.50 | -36.24 |
| 78 | PPVDYWMLLAPTAAG | 1601.88 | 3.80 | - | 20 | -20.84 | -9.80 | -46.60 |
| 79 | TTQNGSYSQYGPLQS | 1630.67 | 5.18 | - | 7.2 | -7.64 | -1.22 | -48.47 |
| 80 | WKFIDVVKTTQNGSY | 1785.99 | 8.50 | - | 2.8 | -16.69 | -1.21 | -30.01 |
| 81 | APTAAGVVVEGTNNT | 1400.51 | 4.00 | - | 4.4 | -6.20 | -1.00 | -52.29 |
| 82 | VDYWMLLAPTAAGVV | 1605.91 | 3.80 | - | 100 | -22.51 | -9.98 | -39.70 |

Solubility = Poor (-), Good (+)

Table S2. Allergenic assessment of immunogenic peptides

| Method | Assessment |
| --- | --- |
| Mapping Igε | Non allergen* |
| MAST[1] | Non allergen* |
| SVM[2] based on amino acid composition | Allergen[$,#] |
| Blast result | Non allergen* |
| Hybrid approach[3] | Non allergen* |

*The protein sequence does not contain experimentally proven Igε epitopes. [1]MAST = Motif Alignment and Search Tool. [2]SVM = Support vector machines. [3]Hybrid approach includes SVM based on dipeptide composition and IgE based approach.

$ For VP7 Score = 0.363, Threshold = -0.4, Positive predictive value = 74.81%, Negative predictive value = 76.94%.

### For VP8 Score =0.134, Threshold, Positive predictive value = 70.05%, Negative predictive value = 80.74%.

Table S3. Population coverage percentage offered by the best predicted epitopes specific for class-I and II-MHC molecule

| Population | VP7 | | VP8 | |
|---|---|---|---|---|
| | MHC-I | MHC-II | MHC-I | MHC-II |
| South Africa | 1.4 | 17.05 | 1.4 | 22.68 |
| Central America | 7.77 | 50.21 | 9.07 | 63.14 |
